# Dot1L methyltransferase activity is a barrier to acquisition of pluripotency but not transdifferentiation

**DOI:** 10.1101/835413

**Authors:** Coral K. Wille, Rupa Sridharan

**Author notes:** Corresponding author: Dr. Rupa Sridharan, Wisconsin Institute for Discovery, University of Wisconsin, 330 North Orchard Street, Room 2118, Madison, WI 53715.

## Abstract

The ability of pluripotent stem cells to be poised to differentiate into any somatic cell type is partly derived from a unique chromatin structure that is depleted for transcriptional elongation associated epigenetic modifications, primarily H3K79 methylation. Inhibiting the H3K79 methyltransferase, Dot1L, increases the efficiency of reprogramming somatic cells to induced pluripotent stem cells (iPSCs) most potently at the mid-point of the process. Surprisingly, despite the enrichment of H3K79me2 on thousands of actively transcribed genes, Dot1L inhibition (Dot1Li) results in few changes in steady state mRNA levels during reprogramming. Dot1Li spuriously upregulates genes not involved in pluripotency and does not shutdown the somatic program. Depletion of the few genes that are downregulated, such as *Nfix*, enhances reprogramming efficiency in cooperation with Dot1Li. Contrary to the prevalent view, Dot1Li promotes iPSC generation beyond early phases of reprogramming such as the mesenchymal to epithelial transition and from already epithelial cell types including keratinocytes. Significantly, Dot1L inhibition does not enhance lineage conversion to neurons or muscle cells. Taken together, our results indicate that H3K79me is not a universal barrier of cell fate transitions but specifically protects somatic cells from reverting to the pluripotent state.

## INTRODUCTION

Differential gene expression allows for functional diversity that translates to tissue specialization in multicellular organisms. In response to signaling and spatial cues, transcription factors engage with the epigenome to culminate in gene expression patterns that establish cell identity (Kelsey *et al*, 2017). The abundance of specific epigenetic modifications changes dynamically during development, right from the formation of the zygote (Dang-Nguyen & Torres-Padilla, 2015; Smith & Meissner, 2013). Fertilization of an oocyte triggers a decrease in Histone 3 Lysine(K) 79 dimethylation (H3K79me2) independent of genome replication (Ooga *et al*, 2008). H3K79me2 levels remain low during the pre-implantation phase until the blastocyst stage (Ooga *et al*, 2008). H3K79me2 is the most differential histone modification between somatic cells and embryonic stem cells (ESCs) that are derived from the blastocyst (Sridharan *et al*, 2013).

Dot1L (disruptor of telomeric silencing-like 1 like) is the sole methyltransferase that performs all levels of H3K79 methylation (me1, me2 and me3) (Black *et al*, 2012). Dot1L knock-out (KO) mice are embryonic lethal between days 9.5 and 13.5 demonstrating the importance of H3K79me in development (Feng *et al*, 2010; Jones *et al*, 2008). Dot1L KO results in disorganized yolk sacs containing primitive erythrocytes. Dot1L is also necessary for development of the heart (Nguyen & Zhang, 2011), cerebral cortex (Franz *et al*, 2019), chondrocytes (Castaño Betancourt *et al*, 2012), and normal CD8+ T cell differentiation (Kwesi-Maliepaard *et al*, 2020).

ESCs isolated from Dot1L KO blastocysts maintain pluripotency, but have decreased proliferation and increased apoptosis (Jones *et al*, 2008). This could be due to defects in chromosomal partitioning as Dot1L KO cells were frequently aneuploid (Jones *et al*, 2008). In contrast, ESCs depleted for Dot1L using shRNA do not have proliferation defects; however, upon differentiation with retinoic acid, cells become aneuploid and arrest in the G2/M phase of the cell cycle (Barry *et al*, 2009). Thus, pluripotent stem cells may have an enhanced ability to tolerate low levels of H3K79me relative to more differentiated cells.

H3K79me2 is quickly demethylated in the reprogrammed nucleus during somatic cell nuclear transfer upon parthenogenesis (Ooga *et al*, 2008). We and others have shown that Dot1L is a barrier for transcription factor mediated reprogramming to induced pluripotent stem cells (iPSCs) from mouse neural stem cells and human fibroblasts (Jackson *et al*, 2016; Onder *et al*, 2012).

Collectively, these phenotypes provide evidence for the importance of Dot1L in cell fate determination; however, the function of H3K79me in mammals has remained elusive. H3K79me2 is enriched on the bodies of rapidly elongating genes (Duffy *et al*, 2018; Veloso *et al*, 2014) implicating the modification as a positive regulator of transcription. In acute leukemia, fusion proteins between mixed-lineage leukemia (MLL) and numerous Dot1L associated proteins (ENL, AF9, and AF10) frequently drive oncogenesis (Mohan *et al*, 2010). MLL target genes such as the HOXA cluster are upregulated concurrent with the corresponding locus becoming H3K79 methylated as Dot1L is mislocalized via the MLL-Dot1L interacting fusion protein. In contrast, H3K79me enrichment directly downregulates the expression of the epithelial sodium channel gene in mouse cells (Zhang *et al*, 2006), confounding the role of Dot1L in transcription.

Here we use the dynamic system of somatic cell reprogramming from mouse embryonic fibroblasts (MEFs) to iPSCs to investigate Dot1L function in maintaining cell identity. We find that Dot1L inhibition enhances MEF pluripotency acquisition throughout the process but acts most potently at mid-reprogramming. This dramatic increase in pluripotency acquisition is accompanied by few transcriptional changes, the majority of which are spuriously upregulated lineage genes that are not enriched for H3K79me2 in MEFs. Modulation of Dot1L regulated genes cannot substitute for chemical inhibition suggesting that loss of H3K79me does not increase reprogramming through single gene transcriptional alterations. We find that one of the Dot1L regulated genes, *Nfix*, maintains cellular identity collaboratively with Dot1L. Dot1L inhibition increases expression of the epithelial marker CDH1 and enhances MEF reprogramming post upregulation of the epithelial program. Epithelial somatic cells such as keratinocytes also display an increased level of H3K79me2, and Dot1L inhibition increases their reprogramming. Surprisingly, Dot1L inhibition decreases MEF lineage conversion into neurons, and does not affect conversion to muscle cells, demonstrating that H3K79me reduction does not generally facilitate cell fate transitions. Thus, despite the abundance of H3K79me2 enrichment in highly transcribed genes, low levels of H3K79me promote pluripotency acquisition without causal alterations in steady state mRNA.

## RESULTS

### Dot1L enzymatic activity is a barrier for reprogramming

We used a secondary reprogramming system to assess how Dot1L affects MEF pluripotency acquisition. MEFs isolated from mice that contain OCT4, SOX2, KLF4, and c-MYC (OSKM) (Sridharan *et al*, 2013) under a doxycycline inducible promoter were reprogrammed in the presence or absence of two Dot1L small molecule inhibitors (Fig 1A, S1A). SGC0946, hereafter called Dot1Li, disrupts Dot1L interaction with S-adenosyl methionine (SAM) by inducing a conformational change and is 10-fold more potent than the first generation Dot1Li inhibitor, EPZ004777 (Yu *et al*, 2012). EZP5676, known as pinometostat, is a highly selective Dot1L inhibitor that also prevents SAM interaction and has entered clinical trials (Campbell *et al*, 2017). Both chemical inhibitors decreased H3K79me2 with similar kinetics compared to control (Fig 1B), and increased reprogramming of MEFs that express varying levels of reprogramming factors based on genotype (one or two copies of the reverse tetracycline transactivator) as measured by NANOG expression (Fig 1A, S1A). Importantly, Dot1L inhibition produced at least 9-fold more *bona fide* pluripotent colonies that maintained NANOG expression after removal of transgene expression by withdrawal of doxycycline (Fig 1A, S1A, right). To determine the temporal requirement for Dot1Li to enhance reprogramming, we performed a 6 day timecourse (Fig 1C, left). When reprogramming populations were exposed to Dot1Li in 2 day intervals, the greatest increase in reprogramming was observed with treatment between days 2-4 (Fig 1C, S1B, blue bars). Removing Dot1Li between days 2-4 resulted in fewer NANOG positive colonies compared to 4 days of continuous Dot1Li treatment either early or late in reprogramming (Fig 1C, S1B, red bars). The greatest number of NANOG+ colonies formed when Dot1L was inhibited throughout reprogramming. Thus, Dot1L activity is a barrier throughout reprogramming but has the greatest effect in the intermediate phase.

**Figure 1.**
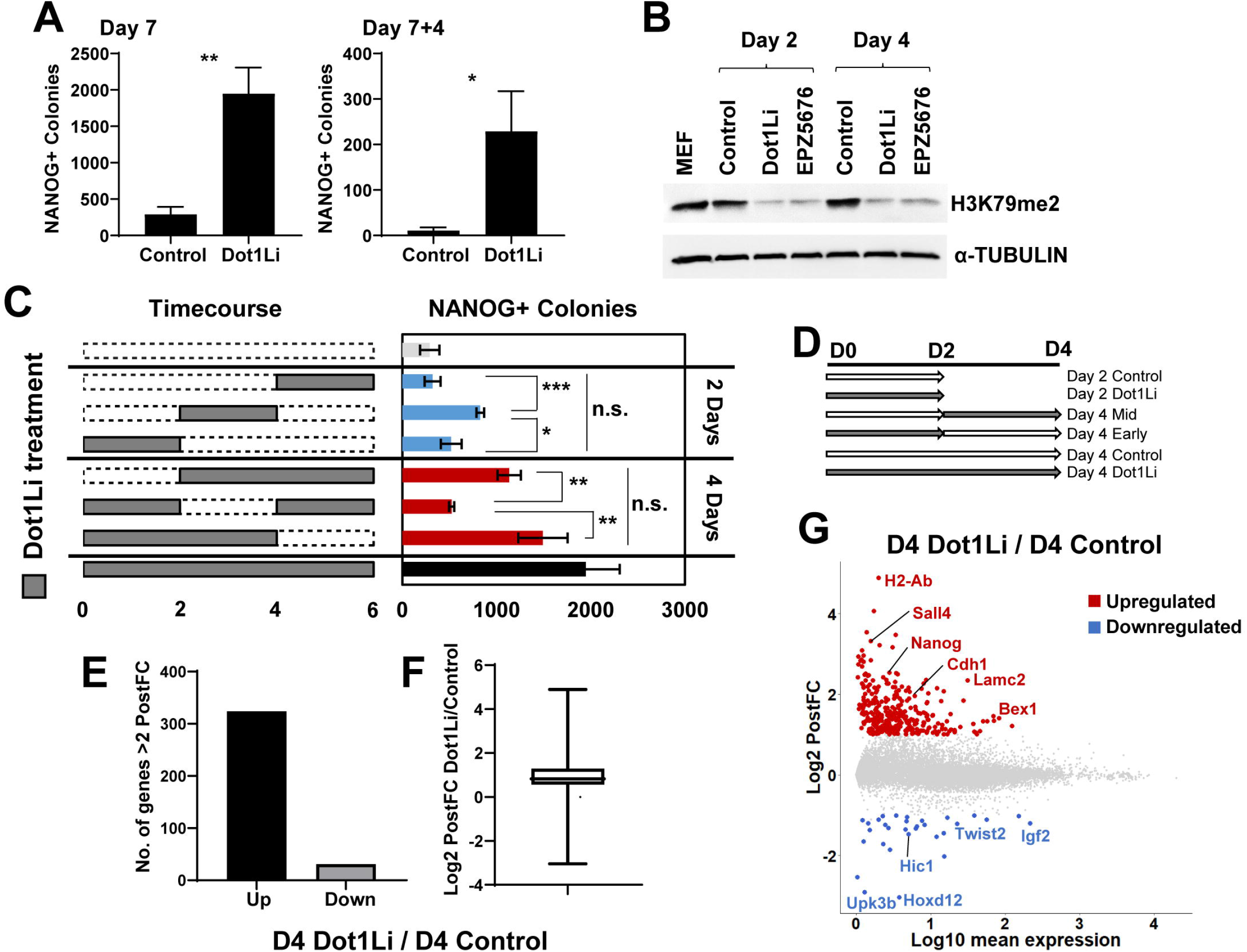
H3K79me is a barrier for MEF reprogramming with few transcriptional effects. A. Number of NANOG+ colonies obtained on day 7 after induction of OSKM in MEFs (Left) and NANOG+/DPPA4+ colonies 4 days post-dox and drug removal (Right) with control DMSO or Dot1L inhibitor SGC0946 (called Dot1Li henceforth). Three biological replicates, each consisting of at least two technical replicates, were assessed. Error bars represent the standard deviation of three technical replicates of a representative biological replicate. **P<0.01 and *P<0.05 by unpaired t□test. B. Immunoblot of H3K79me2 and TUBULIN loading control of reprogramming cells on day 2 or 4 treated with control, Dot1Li, or EPZ-5676. C. Left panel: Scheme of exposure to Dot1Li (gray boxes) or control (dotted boxes). Right panel: Number of NANOG+ colonies on day 6. Four biological replicates, each consisting of at least two technical replicates, were assessed. Error bars represent the standard deviation of three technical replicates of a representative biological replicate. ***P<0.001, **P<0.01, *P<0.05 by unpaired t□test. D. Schematic of RNA-Seq timepoints. E. Number of genes upregulated or downregulated more than 2-fold (PostFC) with a posterior probability of differential expression greater than 0.95 determined by EBSeq in day 4 Dot1Li versus day 4 Control. F. Box plot of Log2 PostFC of all genes with a posterior probability of differential expression greater than 0.95 in day 4 Dot1Li relative to day 4 Control. G. Log10 average TPM of day 4 Dot1Li and 4 Control versus Log2 PostFC of all genes. More than 2-fold upregulated indicated in red and downregulated indicated in blue.

### Loss of H3K79me during reprogramming results in few transcriptional changes

To elucidate how loss of H3K79me enhances iPSC generation, the starting MEFs, reprogramming populations on days 2 and 4 days (Fig 1D), as well as ESCs were transcriptionally profiled. Cells collected on day 4 were treated continuously with Dot1Li, treated “early” from days 0-2, treated “mid” from days 2-4, or treated continuously with vehicle control to capture how Dot1Li specifically increases mid reprogramming (Fig 1C, S1B). Continuous Dot1L inhibition for four days resulted in the most transcriptional alterations compared to Early or Mid treatment (Fig 1E, S1C-E). 352 genes were differentially expressed (Fig 1E, Methods) with ten-fold more genes upregulated than downregulated. The majority of differentially expressed (DE) genes had a modest change in expression (Fig 1F), of genes with low levels of absolute expression (Fig 1G). Mid treatment from days 2-4 with Dot1Li yielded the next highest number of changes with 125 upregulated genes and 2 downregulated genes (Fig S1E), followed by day 2 Dot1Li (51 upregulated and 1 down) (Fig S1C). Removal of Dot1Li after 2 days (Early treatment) resulted in only 9 DE genes (Fig S1D). Thus, Dot1L inhibition first results in gene upregulation followed by downregulation of select genes, and these expression changes require sustained inhibition. At every timepoint, a far greater number of upregulated than downregulated genes were DE.

### H3K79me2 is enriched on numerous genes yet few change transcriptionally

Dot1L inhibition could promote iPSC generation by erasing the somatic program, enhancing pluripotent gene expression, or a combination of the two. To distinguish between these possibilities, significantly DE genes from all timepoints (n=438, “Dot1Li-DE”) were compared to the change in gene expression in ESCs versus MEFs. While about 50% of Dot1Li downregulated genes were also decreased in ESCs, far fewer of the Dot1Li upregulated genes were upregulated in ESCs; suggesting that most of the transcriptional change is aberrant to that observed in ESCs. (Fig 2A).

**Figure 2.**
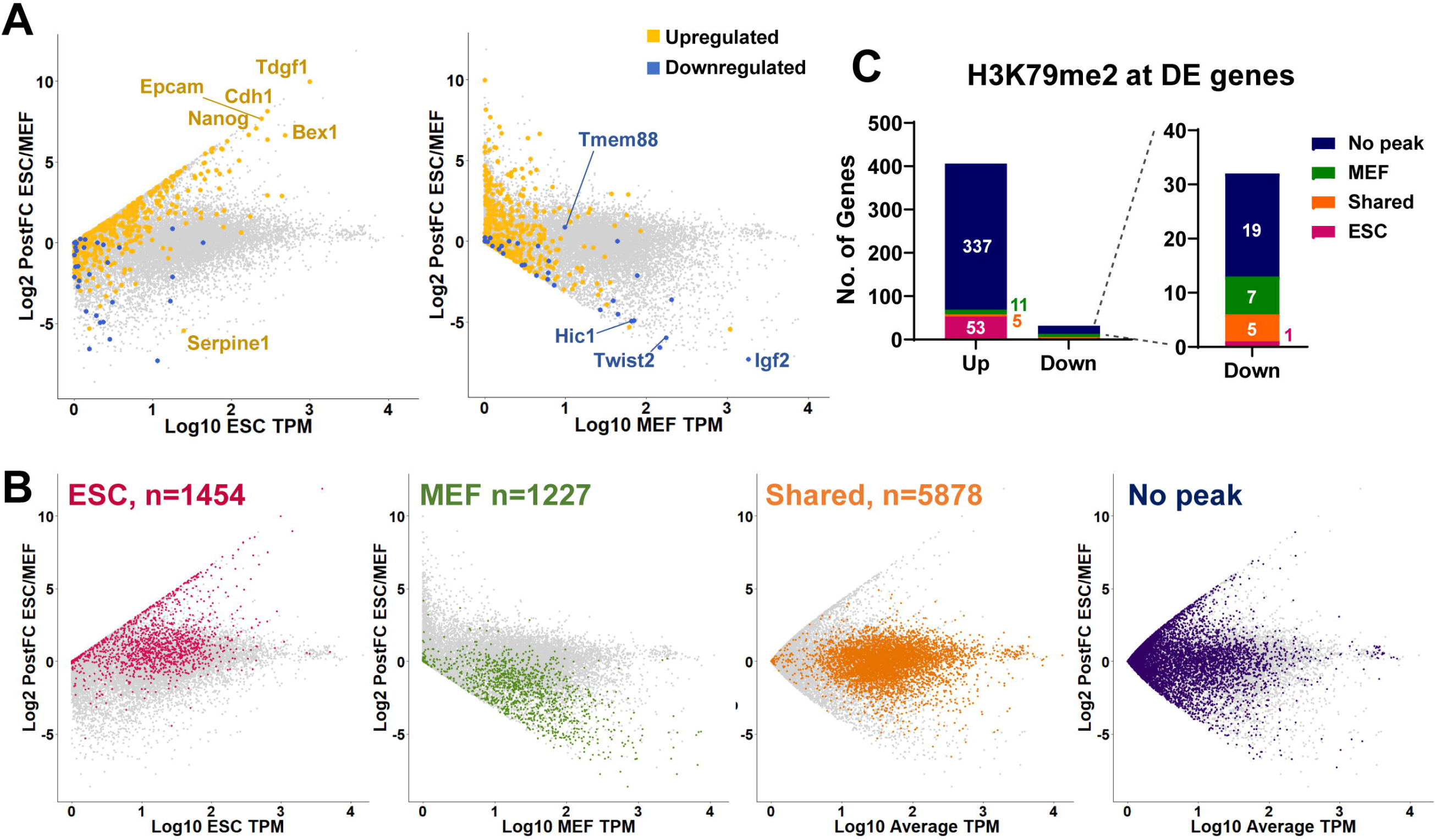
H3K79me2 is enriched on numerous genes yet few change transcriptionally. A. Genes upregulated (gold) or downregulated (blue) by Dot1Li treatment at any time point relative to the matched control (Dot1Li-DE list) plotted on Log10 ESC (Left) or MEF (Right) TPM versus Log2 PostFC in ESCs relative to MEFs. B. Genes with H3K79me2 called peaks in ESCs (red), MEFs (green), shared in both cell types (orange), or with no peak (purple) were plotted on Log10 of the indicated TPM versus Log2 PostFC in ESCs relative to MEFs. H3K79me2 location data were analyzed from GEO GSE90895 (Chronis *et al*, 2017). C. Bar graph of H3K79me2 called peaks at Dot1Li-DE genes.

We then asked whether the transcriptional changes were correlated with the presence of H3K79 methylation. ChIP-Seq of H3K79me2 in MEFs derived from the same reprogrammable mouse line and ESCs was analyzed (Fig 2B) (Chronis *et al*, 2017). As expected, highly expressed genes were enriched for H3K79me2 in both MEFs and ESCs. Genes with “shared peaks” were highly expressed and predominantly housekeeping genes. Additionally, many lowly expressed genes did not have a significant H3K79me2 peak, yet a subset of these genes were DE in ESCs vs. MEFs (Fig 2B, “No peak”).

Although there are 7,105 genes with H3K79me2 in MEFs, only 438 genes are significantly DE upon Dot1L inhibition across all timepoints. The majority of DE genes did not have an H3K79me2 peak (Fig 2C, S2). Thirteen percent of upregulated genes had a peak in ESCs while only 2.7% had a peak in MEFs, and 1.2% had a shared peak (Fig 2C). Only 22% of downregulated genes have a peak in MEFs, 16% have a shared peak, and 1 gene had a peak in ESCs (Fig 2C). Therefore, Dot1Li mediated transcriptional regulation is largely indirect and not coincident only with the genes that contain H3K79me2.

### Dot1L inhibition promotes aberrant transcriptional changes

To ascertain the functional role of Dot1L inhibition during reprogramming irrespective of timepoint, all Dot1Li-DE genes were clustered and visualized on a heatmap relative to their expression in MEFs (Fig 3A). Genes in clusters I-III were changed by Dot1Li to resemble ESC-like expression and included transcription factors involved in structure morphogenesis, epithelium, and proliferation genes (Fig 3A–C).

**Figure 3.**
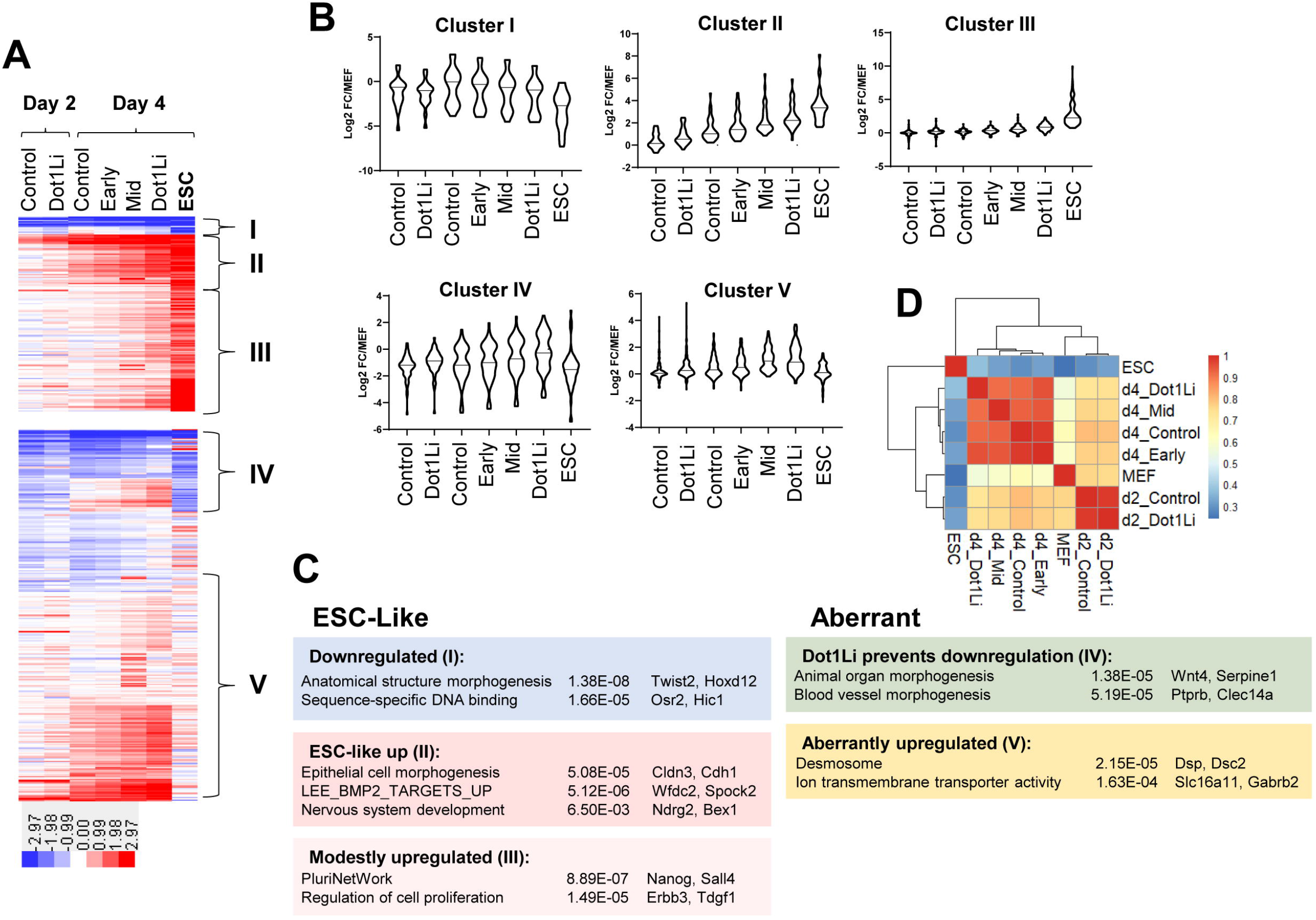
Dot1L inhibition promotes aberrant transcriptional changes. A. k-means clustering of Dot1Li-DE genes organized based on transcriptional change in ESCs relative to MEFs. Heatmap intensity indicates Log2FC TPM relative to MEF for each sample. B. Violin plot of Log2FC TPM relative to MEFs for each cluster. C. Representative gene ontology categories for each cluster. The most enriched categories containing unique sets of genes are displayed (HOMER). D. Spearman correlation of Dot1Li-DE gene TPM across all samples.

Although the changes in clusters I-III (Fig 3C) position Dot1Li treated cells closer to pluripotency acquisition than control treated cells, Spearman correlation of Dot1Li-DE gene expression shows that all timepoints more closely resemble MEFs than ESCs (Fig 3D). Both day 2 samples cluster together and are more correlated to day 4 Control compared to other day 4 samples, suggesting Day 4 Control treatment is kinetically behind the other Dot1Li treatments. Day 4 Early (d0-2) clusters with day 4 Control explaining why a removal of Dot1Li on day 2 of reprogramming barely increases NANOG+ colony formation relative to control treatment (Fig 1C). There was a boost in reprogramming of cells treated from days 2 to 4 with Dot1Li (Fig 1C), yet interestingly, the transcriptional profile of Day 4 Mid (d2-4) clusters closest to day 4 Early (d0-2). Thus, Dot1Li mediated transcriptional effects may not immediately follow reduction of H3K79me2.

By contrast to genes in clusters I-III, the majority of genes (clusters IV and V) do not resemble the transcriptional profile of ESCs relative to MEFs (Fig 3A). Genes in cluster IV (n=60) are either aberrantly upregulated or their downregulation is prevented by Dot1Li (Fig 3A–B) and are not functionally related. Cluster V (n=160) contains genes that are upregulated by Dot1Li treatment yet are largely unchanged in ESCs relative to MEFs (Fig 3A–B) that function in transmembrane signaling (Fig 3C).

To determine how the clusters may be regulated, we performed motif analysis and found that cluster III, and to a lesser extent II, were enriched for E-boxes that can be bound by a cadre of proteins depending on the motif context (Fig S3A). Mesenchymal E-box binding genes that repress expression are not differentially expressed by Dot1Li, with the exception of *Twist2*, which is 2-fold downregulated on day 4 with constant Dot1Li inhibition (Fig S3B). Of note, Dot1Li inhibition also activates *Mycn* (2.7-fold in day 4 Dot1Li) that binds E-boxes to promote gene activity. Thus, the balance of these two proteins may be responsible for the modest increase in expression of cluster III, which is much more significantly enriched for E-boxes compared to cluster II (Fig S3A). Cluster IV is enriched for TCF7 motifs, a transcriptional activator, which is upregulated by Dot1Li treatment. Finally, cluster V is enriched for HOXD12 and HIC1 motifs, both transcriptional regulators downregulated by Dot1Li treatment (Fig 4D).

**Figure 4.**
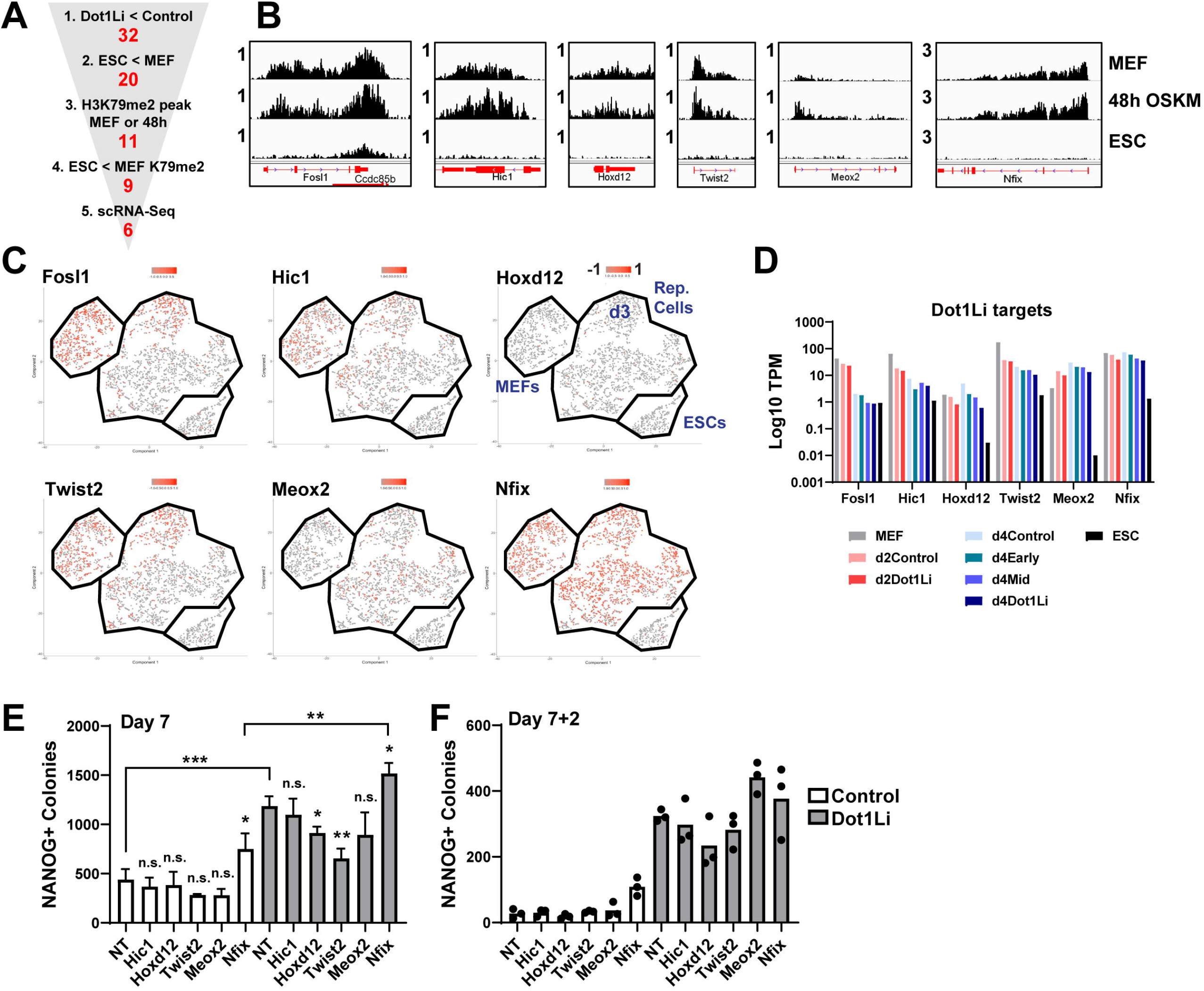
Inhibition of Dot1L enhances reprogramming beyond modulation of single genes. A. Schematic to identify Dot1L direct target genes. Genes were filtered for: 1) At least 2-PostFC downregulation in expression by Dot1Li treatment at any timepoint, 2) At least 2-PostFC downregulation in expression in ESC vs. MEFs, 3) A significant genebody H3K79me2 peak (p-4) with 3-fold enrichment over input in MEFs or 48h OSKM (Fig 4B), 4) A 2-fold reduction in H3K79me2 genebody reads in ESCs, and 5) Expression in a significant portion of MEFs or reprogramming cells by single cell RNA-seq (Fig 4C). B. IGV tracks of H3K79me2 signal in MEFs, 48hs OSKM, and ESCs of potential Dot1L direct targets. C. Cells expressing the Dot1L target gene are labeled in red on the scRNA-seq t-SNE. MEFs, reprogramming cells (days 3, 6, 9, and 12), and ESCs are indicated and labeled on right. Day 3 of reprogramming is indicated but the other days are mixed as reprogramming becomes more heterogenous as it proceeds (Tran *et al*, 2019). D. Log10 TPM bargraph of candidate Dot1Li downregulated genes. E. Number of NANOG+ colonies on day 7 of reprogramming of cells transfected with nontargeting (NT) and siRNA against the indicated Dot1L direct target gene, treated with control (white bars) or Dot1Li (gray bars). Three biological replicates, each consisting of three technical replicates, were assessed. Error bars represent the standard deviation of three technical replicates of a representative biological replicate. ***P<0.001, **P<0.01, *P<0.05, and not significant (n.s.) P>0.05 by unpaired t□test. F. Number of transgene independent colonies two days post dox removal (7+2) of cells transfected with nontargeting (NT) and siRNA against the indicated Dot1L direct target gene, treated with control (white bars) or Dot1Li (gray bars). Two biological replicates, each consisting of three technical replicates, were assessed. Each technical replicate of a representative biological replicate is depicted as a black dot.

Thus, Dot1Li may promote development of reprogramming intermediates expressing lineage genes (Polo *et al*, 2012) due to subtle changes in transcription factor levels. However, as the two clusters that most resemble ESCs (I and II) are devoid of motifs bound significantly by any of the Dot1Li DE genes, it is unlikely that Dot1L regulates pluripotency acquisition through such intermediates.

### Inhibition of Dot1L enhances reprogramming beyond modulation of single genes

Since Dot1Li did not change the entire transcriptional landscape either away from a somatic program or toward a pluripotency program, we used a candidate approach to screen Dot1Li-DE genes for their contribution to pluripotency acquisition. As a higher percentage of downregulated genes were modified by H3K79me2 (Fig 2C, S3C), and Dot1L has previously been reported as a barrier of reprogramming by maintaining fibroblast gene transcription (Onder *et al*, 2012), we comprehensively ranked Dot1Li-directly downregulated genes (Fig 4A, step 1). Among the Dot1Li genes that were downregulated in ESCs vs. MEFs (step 2), we filtered those that had a significant H3K79me2 peak in MEFs or 48h post-OSKM induction (step 3) that is reduced in ESCs (step 4) (Fig 4B). A change in gene expression levels between MEFs and ESCs could occur in every single cell of the starting population or could reflect a population average. We have previously performed single cell RNA-Seq of MEFs, ESCs and a time course of reprogramming (Tran *et al*, 2019). Using this data, we then chose genes that were expressed in the entire population of starting MEFs or reprogramming cells (step 5) (Fig 4C) narrowing to six potential targets that could be critical for Dot1L inhibition (Fig 4D).

Each target was reduced individually using siRNA-mediated knockdown in the presence or absence of Dot1Li. Depletion of *Fosl1* resulted in cell death with three different siRNAs (data not shown) and was not analyzed further. For each of the other five targets, siRNA mediated depletion was robust and resulted in a similar fold change as that obtained by Dot1Li (Fig S4A). When combined with Dot1Li, mRNA levels were further reduced (Fig S4A). *Hic1*, *Hoxd12*, and *Twist2* depletion did not affect reprogramming in the presence or absence of Dot1Li (Fig 4E–F, S4B-C). *Meox2* depletion did not increase NANOG+ colony formation during OKSM transgene expression, but enhanced *bona fide* colony formation in combination with Dot1Li (Fig 4E–F, S4B-C). Depletion of *Nfix* increased the occurrence of NANOG+ colonies 1.6-fold compared to a 2.5-fold increase with Dot1Li (Fig 4E, S4B). However, *Nfix* depletion did not lead to a robust formation of *bona fide* colonies as Dot1Li treatment increased transgene independent colonies 11-fold whereas only a 4.7-fold increase was observed with *siNfix* (Fig 4F, S4C). When combined with Dot1Li, *Nfix* depletion enhanced both NANOG+ and transgene independent colony formation. Thus, the downregulation of *Nfix* and *Meox2* act in an additive manner with Dot1Li to increase pluripotency by contributing to the Dot1Li phenotype.

By comparing the results of the depletion experiments (Fig 4E–F) with the pattern of expression from single cell analysis (Fig 4C) we find that Dot1Li acts in a collaborative manner with genes that are still present at later stages of reprogramming. *Hic1*, *Fosl1* and *Twist2* are downregulated in most cells by day 3 of reprogramming. *Meox2* is upregulated only in reprogramming cells and *Nfix* continues to be expressed in most reprogramming cells. Taken together with the temporal effectiveness of Dot1Li at the mid stages of reprogramming (Fig 1C), these data suggest that Dot1Li sets the stage for enhancing reprogramming efficiency along with downregulation of persistently expressed genes such as *Meox* and *Nfix*. Furthermore, *Nfix* collaboratively enhances Dot1Li-mediated colony formation and *Meox2* specifically enhances a later stage of reprogramming suggesting they function in separate pathways.

### Inhibition of Dot1L enhances reprogramming of epithelial cells

To investigate if single genes in the upregulated cohort could replace Dot1Li, we examined the epithelial genes. Mesenchymal to epithelial transition is an early phase of reprogramming when starting with MEFs (Apostolou & Hochedlinger, 2013). Corroborating a previous report on Dot1Li enhancing epithelial expression (Onder *et al*, 2012), Cluster II genes (Fig 3A) that were most similar in their expression changes to ESCs (Fig 3B) are enriched for epithelial genes. We introduced *Cdh1*, which has previously been shown to be required for pluripotency (Chen *et al*, 2010; Hawkins *et al*, 2012), in reprogrammable MEFs using a lentivirus to achieve expression level comparable to that in ESCs (Fig 5A). Although CDH1 was detected at the cell surface (Fig 5A), it did not enhance MEF reprogramming (Fig 5B). It is important to note that exogenous *Cdh1* expression was incapable of causing mesenchymal gene downregulation (Fig S5A). This result supports the notion that mesenchymal gene downregulation and *Cdh1* upregulation are regulated independently of each other from our co-expression analysis of scRNA-Seq data, where we found that reprogramming cells can simultaneously express genes from both programs (Tran *et al*, 2019). Given that expression of *Cdh1* alone could not replace Dot1Li function (Fig 5B), we assessed how reduction of H3K79me affects reprogramming of cells that are already epithelial. Reprogramming MEFs were sorted on day 3.5 post OSKM induction for cell surface CDH1 expression (Fig 5C) to a similar level as ESCs (Fig S5B). Reprogramming of the CDH1- and CDH1+ sorted cells was then continued in the presence of Dot1Li. Interestingly, reprogramming of both populations of cells was increased to a similar extent (4-fold) when exposed to Dot1Li (Fig 5C, S5C) indicating a response beyond the upregulation of the epithelial program.

**Figure 5.**
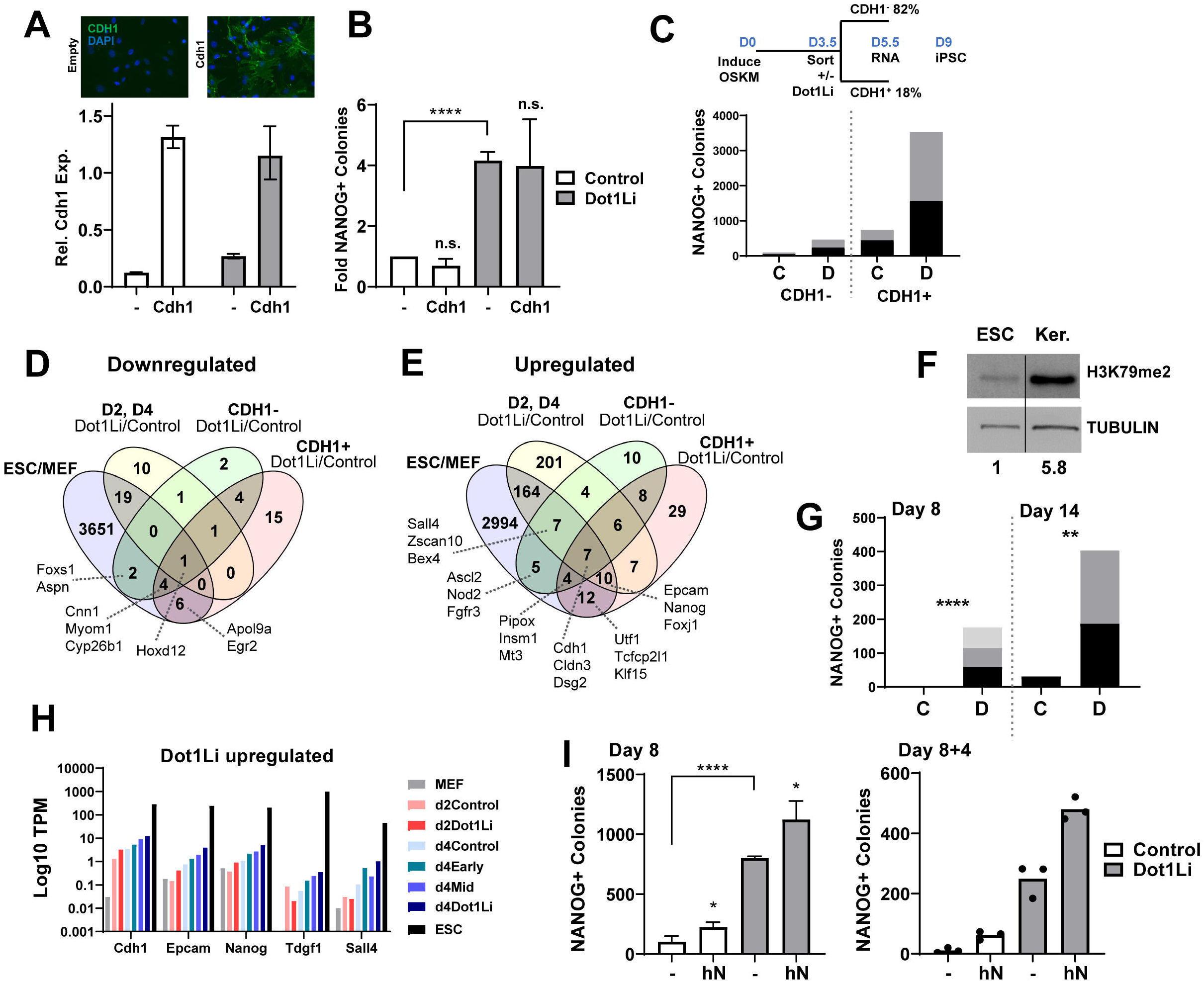
Inhibition of Dot1L enhances reprogramming of epithelial cells. A. Top: CDH1 immunofluorescence of day 4 of reprogramming of empty vector (-) and *Cdh1* transduced cells. Bottom: Relative *Cdh1* expression measured on day 3 of reprogramming, ESCs set to 1. Cells were treated with control (white bars) or Dot1Li (gray bars). B. Fold NANOG+ colonies of empty vector (-) or *Cdh1* transduced cells, treated with control (white bars) or Dot1Li (gray bars). Empty vector transduced cells treated with control set to 1. Error bars represent the standard deviation of three biological replicates each consisting of two technical replicates. ****P<0.0001 and not significant (n.s.) P>0.05 by unpaired t test. C. Reprogramming scheme (Top): Cells were grown for 3 days in KSR containing media to accelerate CDH1 expression. On day 3.5, cells were sorted with flow cytometry for CDH1 surface expression, followed by Dot1Li or control treatment. RNA-Seq samples were collected at day 5.5 and reprogramming was evaluated on day 9. Bottom: NANOG+ colony formation of CDH1 sorted cells treated with Dot1Li (D) or control (C). Two biological replicates, each consisting of two technical replicates, were assessed. Each technical replicate of a representative biological replicate is depicted as stacked bars. D. Overlap of 1.5 PostFC Dot1Li downregulated genes in CDH1- and CDH1+ cells with the downregulated genes from all timepoints of the Dot1Li timecourse (2 PostFC) (Fig 3A) and genes downregulated in ESCs relative to MEFs (2 PostFC). A selection of genes is displayed. E. Overlap of 1.5 PostFC Dot1Li upregulated genes in CDH1- and CDH1+ cells with the upregulated genes from all timepoints of the Dot1Li timecourse (2 PostFC) (Fig 3A) and genes upregulated in ESCs relative to MEFs (2 PostFC). A selection of genes is displayed. F. Immunoblot of H3K79me2 and TUBULIN loading control in ESCs and keratinocytes (Ker.). G. NANOG+ colonies on day 8 and day 14 of keratinocytes treated with Dot1Li (D) or control (C). Different shades of gray depict 3 technical replicates. ****P<0.0001 and **P<0.01 by unpaired t test. H. Log10 TPM bargraph of Dot1Li upregulated reprogramming factors. I. Cells transduced with vector control (-) or human Nanog (hN), treated with control (white bars) or Dot1Li (gray bars). Left: Number of NANOG+ colonies on day 8 of reprogramming. Two biological replicates, each consisting of three technical replicates, were assessed. Error bars represent the standard deviation of three technical replicates of a representative biological replicate. ****P<0.0001 and *P<0.05 by unpaired tLtest. Right: Number of transgene independent colonies 4 days post dox removal (8+4). Two biological replicates, each consisting of three technical replicates, were assessed. Each technical replicate of a representative biological replicate is depicted as a black dot.

To identify a transcriptional response after the epithelial upregulation, we profiled the gene expression of CDH1- and CDH+ populations with Dot1Li. Similar to the results from the unsorted timecourse (Figs 1, 3), more genes were upregulated than downregulated by Dot1Li in both CDH1- and CDH1+ populations. *Hoxd12* was the only commonly downregulated gene in CDH1-, CDH1+, Dot1Li-DE days 2 and 4, and in ESCs relative to MEFs (Fig 5D). Upregulated genes that were altered in the same direction as their expression in ESCs (Fig 5E) were enriched for the functional categories of stem cell population maintenance and epithelial cell development. However, the upregulation was not to the same extent as ESCs (Fig S5D), as previously observed (Fig 3).

Another strategy to assess if Dot1L inhibition affects pluripotency acquisition after cell fate transition is reprogramming of pre-iPSCs. Pre-iPSCs are stalled intermediates of reprogramming that have downregulated the somatic program but have not activated the pluripotency gene regulatory network, yet can be clonally propagated indefinitely similar to iPSCs (Silva *et al*, 2008; Sridharan *et al*, 2009). In a MEF-derived pre-iPSC line with high expression of CDH1 (Fig S5E), Dot1Li treatment greatly reduced H3K79me2 levels (Fig S5F). Pre-iPSCs can be converted to iPSCs at low efficiency by exposure to ascorbic acid (AA) (Tran *et al*, 2015). In these pre-iPSCs, Dot1Li treatment alone increased surface CDH1 1.2-fold but did not result in NANOG positive cells (Fig S5G-H). AA alone increased CDH1 and NANOG expression compared to control treated cells. However, Dot1Li treatment in addition to AA increased NANOG expression 2.4-fold without altering CDH1 expression (Fig S5G-H). Thus, CDH1 expression does not directly promote NANOG upregulation, and Dot1Li can increase reprogramming without affecting CDH1 surface expression. Taken together, Dot1L inhibition enhances pluripotency acquisition beyond downregulation of the somatic program and upregulation of the epithelial program.

We next investigated whether Dot1L inhibition could enhance reprogramming from other cell types. Keratinocytes are an epithelial cell type, and despite not having to undergo one of the phases of MEF reprogramming, mesenchymal to epithelial transition (MET) (Li *et al*, 2010; Samavarchi-Tehrani *et al*, 2010), they still reprogram poorly as compared to MEFs (Nefzger *et al*, 2017). We isolated keratinocytes isolated from reprogrammable mice and confirmed that they express similar levels of CDH1 on their surface as ESCs (Fig S5I). Since our mass spectrometry for histone modification enrichment was from MEFs, we performed an immunoblot for H3K79me2. Keratinocytes had over 5-fold more H3K79me2 globally than ESCs (Fig 5F). Dot1Li applied during keratinocyte reprogramming significantly increased the efficiency over 18-fold by day 14 (Fig 5G). The kinetics of reprogramming was also increased with the appearance of the first NANOG+ colonies by day 8 of reprogramming. Thus, reduction of H3K79me2 is not correlated to epithelial identity, rather it promotes pluripotency.

We next assessed the upregulation of pluripotency factors. In a previously reported single cell transcriptomic analysis from our lab, we found that the co-expression of a quartet of genes *Nanog*, *Sall4*, *Tdgf1* and *Epcam* within the same cell predicted a more homogenous transition to an iPSC state (Tran *et al*, 2019). Dot1Li upregulated the same genes but to a much smaller magnitude than that in ESCs (Fig 5H). We introduced exogenous NANOG (Fig S6A-B) and found that it modestly enhanced reprogramming and transgene independence, in both the presence and absence of Dot1Li (Fig 5I, S6C). Thus, Dot1Li does not increase reprogramming through *Nanog* upregulation but collaborates with NANOG especially to maintain pluripotent colonies.

To identify if any other Dot1Li DE genes regulate pluripotency acquisition, they were overlapped with genes identified in a large shRNA screen starting with the same OSKM inducible secondary MEF system (Borkent *et al*, 2016). Among the genes that affected reprogramming more than 2-fold from the screen (~2300 genes), only 10 overlapped. However, 7 barrier genes were upregulated and 3 enhancers were downregulated by Dot1Li, in the opposite direction of pluripotency promotion (Fig S6D). Thus, Dot1L inhibition likely does not increase reprogramming by modulating single gene expression. Additionally, Dot1Li treatment does not increase cellular proliferation (Fig S6E) or cause changes in the cell cycle (Fig S6F-G) indicating it does not enhance pluripotency by increased cell number.

### Depletion of H3K79me does not universally promote cell fate transitions

Given our results that Dot1Li enhanced iPSC generation from somatic cells of different germ layers but with a general lack of a definitive transcriptional response, we wondered if H3K79 methylation was a barrier to all cell fate transitions. If the role of H3K79me2 were to maintain high levels of cell fate specific gene expression, then inhibition of Dot1L should enhance any cell fate transition. Cell fate can be directly altered without going through a pluripotent intermediate by the process of transdifferentiation where transcription factors of the endpoint cell type are expressed in the recipient somatic cell (Davis *et al*, 1987). MEFs can be converted into induced neurons (iN) by the expression of *Brn2*, *Ascl1*, and *Myt1l* (BAM) (Fig 6H) (Vierbuchen *et al*, 2010). MEFs transduced with the BAM transcription factors were monitored for iN formation by class III beta-tubulin (TUJ1) expression, a marker of neuronal cells (Fig 6A). In contrast to the effect on reprogramming to iPSCs, transdifferentiation of MEFs to iNs was not enhanced, but instead reduced with exposure to Dot1Li (Fig 6B, S6H). Expression of neuronal markers Olig2 and Map2 was also decreased upon Dot1Li (Fig 6C).

**Figure 6.**
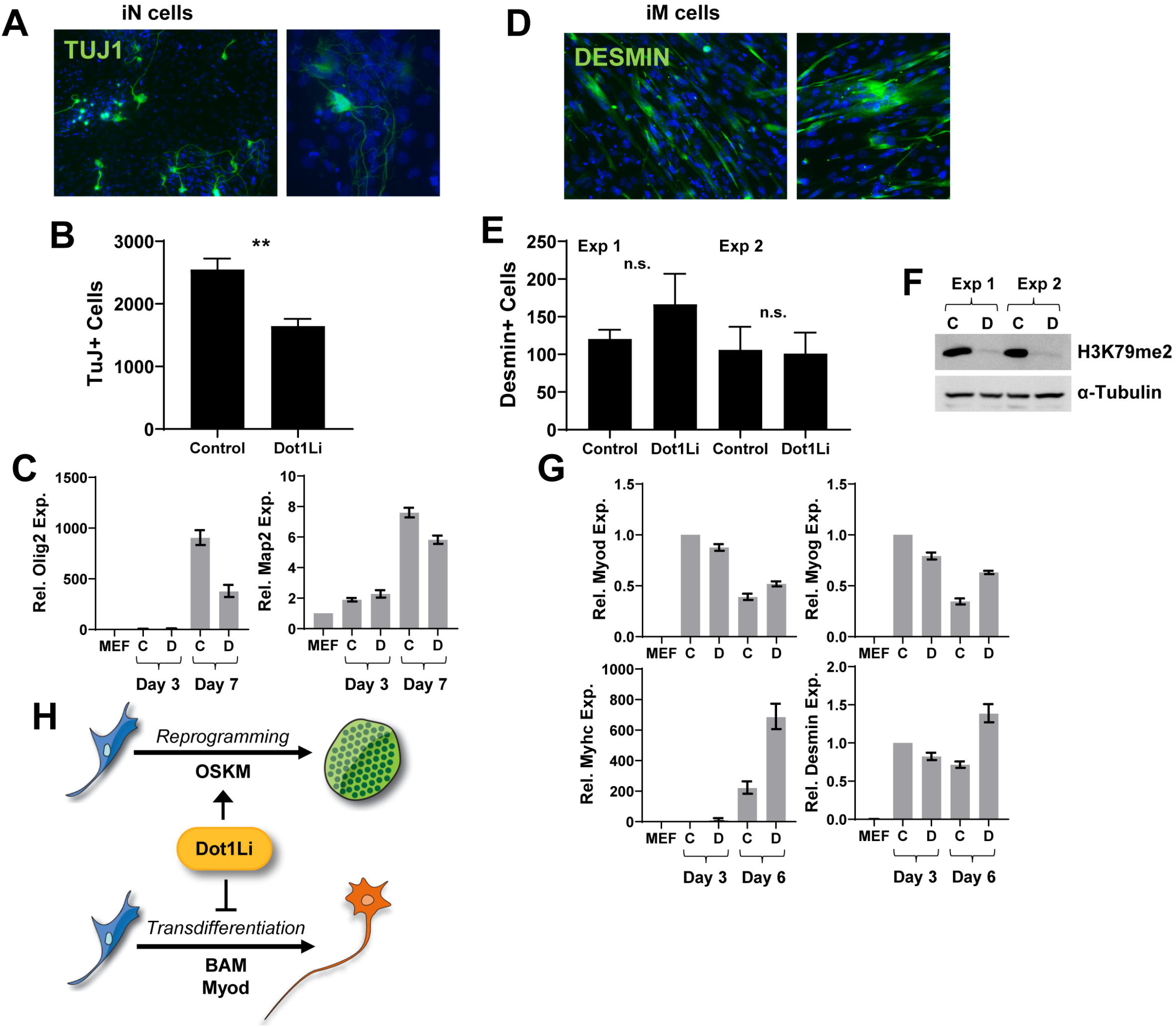
Dot1Li does not universally promote cell fate transitions. A. Representative images of TUJ+ induced neurons (iN). B. TUJ+ iN on day 6 post *Brn2*, *Ascl1*, and *Myt1l* (BAM) induction, in the presence and absence of Dot1Li. Two biological replicates, each consisting of at least two technical replicates, were assessed. Error bars represent the standard deviation of three technical replicates of a representative biological replicate. **P<0.01 by unpaired t test. C. Relative expression of neuronal factors in MEFs and on days 3 and 7 of transdifferentiation treated with control (C) or Dot1Li (D). MEFs set to 1. D. Representative images of DESMIN+ induced muscle cells (iM). E. DESMIN+ iM on day 6 post *Myod* transduction, in the presence and absence of Dot1Li. Error bars represent the standard deviation of three technical replicates of two biological replicates (Exp 1 and 2). Not significant (n.s.) indicates P>0.05 by unpaired t test. F. H3K79me2 immunoblot on day 4 of iM transdifferentiation in cells treated with control (C) or Dot1Li (D). G. Relative expression of muscle factors in MEFs and on days 3 and 6 of iM transdifferentiation treated with control (C) or Dot1Li (D). Cells on day 3 treated with control set to 1. H. Schematic of the role of Dot1Li in cell fate change. Dot1Li is a barrier for MEF pluripotency acquisition but not transdifferentiation to neurons or muscle cells. ESCs are depleted for H3K79me compared to somatic cells. MEFs, neurons, and muscle cells are committed cell types that express their somatic program while repressing other programs. Pluripotency is a unique state where cells must express the pluripotency program yet remain poised to express any other lineage program upon the appropriate cues.

To test transdifferentiation to a cell type derived from a different germ layer, MEFs were converted to induced muscle cells (iM) with MYOD expression (Fig 6H) (Bar-Nur *et al*, 2018). Dot1Li treatment (Fig 6F) did not significantly affect iM formation (Fig 6E) measured by DESMIN expression (Fig 6D). Similar levels of muscle markers *Myog*, *Myhc*, and *Desmin* were expressed on day 3 of transdifferentiation, yet Dot1Li maintained *Myod* transgene expression of day 6 (Fig 6G) which was accompanied by increased muscle marker expression at that timepoint. Therefore, Dot1L is not a general barrier for cell fate transitions, rather reduction of H3K79me specifically promotes pluripotency (Fig 6H).

## DISCUSSION

Dot1L is crucial for mammalian development, yet the role of H3K79me in cell fate determination is still unknown. Here we find that Dot1L is a barrier for pluripotency acquisition of MEFs throughout reprogramming but acts most strongly during mid-reprogramming (Fig 1). Stages of reprogramming when starting from MEFs include an early inactivation of the somatic program, an important component of which are the mesenchymal genes (Li *et al*, 2010; Samavarchi-Tehrani *et al*, 2010). Inhibition of Dot1L was reported to enhance human fibroblast reprogramming by facilitating MET (Onder *et al*, 2012). Although we find that inhibition of Dot1L increases *Cdh1* expression in our mouse reprogramming studies, it also enhances reprogramming of keratinocytes that do not have to undergo MET, and MEF reprogramming beyond MET (Fig 5). Additionally, we recently showed that the epithelial and mesenchymal programs are independently regulated in the presence of Dot1Li during reprogramming using single cell RNA-Seq (Tran *et al*, 2019).

In the course of assessing Dot1L transcriptional regulation, we identified contributions of two reprogramming barriers. The direct targets *Nfix* and *Meox2*, maintain cellular identity with Dot1L, but function at different stages of reprogramming. NFIX is transcription factor important for neural (Pekarik & Belmonte, 2008) and muscle development (Pistocchi *et al*, 2013). Additionally NFIX is required to maintain murine hair follicle stem cell enhancer function (Adam *et al*, 2020) and its depletion increases the number of pluripotent colonies (Yang *et al*, 2011) specifically during exogenous factor expression (Fig 4E-F). From our previous analysis of reprogramming with scRNA-seq, we ordered single cells in a trajectory (Tran *et al*, 2019). We found a major branchpoint where cells stalled and did not complete the transition to iPSCs (Tran *et al*, 2019). *Nfix* was found to be a branchpoint gene such that downregulation was required to continue in the reprogramming trajectory suggesting that it may have a specific role in later reprogramming. MEOX2 is a homeobox transcriptional factor important for somite and limb development, and has been shown to increase reprogramming in a previous study (Pfaff *et al*, 2011). Here we find that depletion of *Meox2* increases *bona fide* colony formation in collaboration with Dot1Li. Whether low levels of *Meox2*, which can only be achieved with both siRNA and Dot1L chemical inhibition (Fig S4A), or loss of H3K79me is required for this phenotype is still unknown. Of the identified Dot1L targets, *Nfix* and *Meox2* continue to be expressed in cells later in reprogramming and they gain H3K79me2 after OSKM induction (Fig 4B–C) suggesting Dot1Li may facilitate their downregulation by preventing the gain in H3K79me2 enrichment at later reprogramming timepoints. It is important to note that depletion of these two factors does not reach the reprogramming efficiency of Dot1Li, and the effect is additive with Dot1L inhibition (Fig 4E–F). Thus, downregulation of these factors may contribute but is not solely casual for the Dot1L reprogramming phenotype.

We find that modulation of Dot1L-DE genes does not substitute for Dot1L chemical inhibition (Figs 4–5) suggesting that H3K79me may have a role in reprogramming beyond transcriptional regulation of single genes. During development, only four genes, were differentially expressed in Dot1L KO mouse c-kit+ cells sorted from E10.5 yolk sacs where profound phenotypic alterations in vascular morphology and erythrocyte maturation were observed (Feng *et al*, 2010). These small transcriptional changes in key factors like *Nanog* during reprogramming or *Gata2* during erythrocyte maturation suggest that loss of H3K79me may alter global epigenetic profiles rather than local expression profiles. For example, loss of H3K79me2 allowed for spread of H3K27me3 on downregulated genes in leukemia cells (Deshpande *et al*, 2014). Both upregulated and downregulated genes can be decorated by H3K79me2 before Dot1L depletion in development of the cerebral cortex (Franz *et al*, 2019). H3K79me2/3 is also enriched at certain intronic enhancers in leukemia cell lines and regulates future deposition of H3K27ac (Godfrey *et al*, 2019), a modification associated with active enhancers.

We observed that many more genes are upregulated rather than downregulated by Dot1Li during reprogramming (Fig 1). Dot1L inhibition could result in de-repression as H3K79me forms a negative transcriptional feedback loop *C. elegans* (Cecere *et al*, 2013). Many of the upregulated genes are expressed in other lineages and are not modified by H3K79me2 in MEFs or ESCs (Fig 3, S3). Alternatively, these genes may be regulated indirectly by the binding of Dot1Li-DE transcription factors such as HOXD12 or HIC1. The aberrant expression of lineage specific genes may promote reprogramming, a notion that can be further investigated by single cell transcriptional analysis. Regardless, the boost to reprogramming cells by Dot1L inhibition outweighs the burden of spurious lineage gene expression.

H3K79me is lower in ESCs than somatic cells (Fig 5F) (Sridharan *et al*, 2013) and further H3K79me depletion does not prevent stem cell renewal (Barry *et al*, 2009; Jones *et al*, 2008). Single-cell RNA-Seq revealed that Dot1L is expressed similarly in MEFs and ESCs (Tran *et al*, 2019) suggesting that pluripotency acquisition is regulated at the level of Dot1L activity. Supporting its direct function in multipotency, loss of H3K79me is associated with gains in plasticity in other systems. For example, rare cells breast epithelial cells can differentiate into multiple lineages when cultured *ex vivo* and require global loss of H3K79me2 (Breindel *et al*, 2017). We find that Dot1L inhibition does not enhance MEF transdifferentiation into cells types derived from two different germ layers: neurons (ectoderm) and muscle cells (mesoderm) (Fig 6), demonstrating that low levels of H3K79me do not universally promote cell fate transitions.

## MATERIALS AND METHODS

### Cell isolation and culture

Male and female MEFs were isolated on day E13.5 from embryos that were homozygous for the Oct4-2A-Klf4-2A-IRES-Sox2-2A-c-Myc (OKSM) transgene at the Col1a1 locus and either heterozygous or homozygous for the reverse tetracycline transactivator (rtTA) allele at the Rosa26 locus, as previously described (Tran *et al*, 2019). MEFs were grown in DMEM, 10% FBS, 1x non-essential amino acids, 1x glutamax, 1x penicillin/streptomycin, and 2-Mercaptoethanol (4 μl/500ml). Feeder MEFs were maintained and isolated as above from DR4 mice genetically resistant to geneticin (G418), puromycin, hygromycin, and 6-thioguanine. Feeder cells were irradiated with 9000 rad after 3 passages. ESC V6.5 were grown on feeder MEFs in knock-out DMEM, 15% FBS, 1x non-essential amino acids, 1x glutamax, 1x penicillin/streptomycin, 2-Mercaptoethanol (4 μl/525ml), and leukemia inhibitory factor. Pre-iPSCs were maintained in ESC medium and were generated by clonally picking and propagating MEFs undergoing retroviral reprogramming that failed to upregulate NANOG as previously described (Silva *et al*, 2008; Sridharan *et al*, 2009). MEFs used for pre-iPSC generation were derived from the NGIP mouse line that contains a NANOG-GFP reporter (Tran *et al*, 2015). Keratinocytes were isolated from reprogrammable mice 4 days postnatal as previously described (Li *et al*, 2017) and cultured in EpiLife™ Medium with 60 μM calcium (ThermoFisher MEPI500CA) with the EpiLife™ Defined Growth Supplement (Thermofisher S0125). 293T were acquired from ATCC and grown in DMEM and 10% FBS. Astrocytes were isolated as previously described (Jackson *et al*, 2016) and maintained in DMEM, 1% FBS, 1x non-essential amino acids, 1x glutamax, and 1x penicillin/streptomycin. Mice were maintained according to the UW-Madison IACUC approved protocol.

### Reprogramming experiments

MEFs were seeded at a density of 30,000 to 50,000 cells/12-well and keratinocytes were seeded at a density of 10,000 cells/24 well onto gelatinized coverslips. Feeder MEFs were added at 1/2x confluency and reprogramming was initiated with 2 μg/ml of doxycycline (dox) with either vehicle (DMSO) control, 5 μM SGC0946 (ApexBio A4167), or 3 μM EPZ5676 (MedChem Express HY-15593). ESC media made with FBS or knock-out serum replacement (KSR), as indicated, with fresh dox and chemicals was replaced every two days. Keratinocytes were maintained in keratinocyte media two days post-OSKM induction before changing to ESC media to avoid FBS-induced differentiation. In the case of exogenous gene expression, MEFs were transduced with lentivirus and selected (if possible) before seeding. Pre-iPSCs were seeded onto gelatinized wells at a density of 200,000 cells/6-well. Reprogramming was initiated with 100 μg/ml of ascorbic acid (Sigma A8960) in ESC media as previously described (Tran *et al*, 2015). Pre-iPSCs were passaged at a density of 200,000 cells/6-well on day 4 of reprogramming and pluripotency acquisition was assessed by NANOG-reporter activity on day 10.

To assess transgene independence, reprogramming cells were washed once with ESC media, and ESC media free of doxycycline and drugs was replaced in wells. Sustained NANOG expression was measured 2-4 days post doxycycline removal by immunofluorescence. Experiments were timed for individual MEFs so that cells were exposed sufficiently to OSKM to produce *bona fide* colonies, but not so long that NANOG positive colonies were overcrowded and uncountable in the presence of doxycycline.

### Induced neuron transdifferentiation

MEFs that were homozygous for reverse tetracycline transactivator (rtTA) allele at the Rosa26 locus were transduced with Tet-inducible *Brn2* (Addgene 27151), *Ascl1* (Addgene 27150), and *Myt1l* (Addgene 27152) lentiviruses. MEFs were plated onto a coverslip at a density of 120,000 cells/per 12 well, 2 days post transduction. Transdifferentiation was initiated with 2 μg/ml of doxycycline as previously described (Vierbuchen *et al*, 2010), with control DMSO or 5 μM SGC0946 (ApexBio A4167). Media was replaced day 3 post-induction with N3 media: DMEM/F12, 1x glutamax, 1x penicillin/streptomycin, 5 μg/ml insulin, 10 ng/ml FGF, and 1 x N-2 (ThermoFisher 17502048) with and without SGC0946. Media was replaced every 2 days until coverslips were fixed for immunofluorescence on days 6-7.

### Muscle cell transdifferentiation

MEFs were transduced with murine *Myod* lentivirus and split in MEF media onto coverslips at 100,000 cells per 12-well with control DMSO or 5 μM SGC0946 (ApexBio A4167) 8 hours post-transduction. On day 2, media was changed to iMPC media: 10% FBS, 10% KSR, 1x Glutamax, 1x non-essential amino acids, 1x penicillin/streptomycin, 2-Mercaptoethanol (4 μl/500ml), 10 ng/ul basic FGF, DMEM (Bar-Nur *et al*, 2018). Media and drugs were refreshed every two days until coverslips were fixed on day 6 for immunofluorescence.

### Lentiviral packaging and transduction

Lentiviral transfer vectors were transfected into 293T cells with packaging vector pspax2 (Addgene 12260) and envelop vector vsvg using linear polyethylenimine. Media was changed to MEF media with 20mM HEPES 4 hours post transfection. Virus-containing media was harvested at 48h and 72h, combined, and filtered through a 0.45 μm PVDF filter. Virus containing media was combined with fresh media at a ratio of 1:1 and 10 μg/ml Hexadimethrine Bromide (polybrene) to transduce target cells.

### RNA isolation and library preparation

RNA was isolated from technical replicates using TRIzol. One-fifth the volume of chloroform was added and phases were separated by max centrifugation. The upper layer was isolated, RNA was precipitated with 0.53 volumes of ethanol, and applied to a RNeasy column (Qiagen 74104). RNA was washed with 500 μl RW1 and DNA was digested on the column with DNAse (Qiagen 79254) for 30 min. RNA was washed according to the RNeasy protocol and eluted in 30 ul of H2O. RNA was quantitated and 1 μg of each sample was combined with 20 ng of RNA from 293T (human) cells, prepared as above, as a spike-in control for sequencing normalization (timecourse and CDH1 sort studies). The cDNA library was constructed using TruSeq RNA Sample Preparation kit V2 (RS-122-2002) according to the manufacturer’s instructions. Libraries were assessed with Qubit and Bioanalyzer3.0.

### RNA-Seq computational analysis

Greater than 40 million reads of timecourse Dot1Li libraries (Fig 1) and CDH1 sort libraries (Fig 5) were sequenced PE150 by Novogene on an Illumina HiSeq 4000. Greater than 25 million reads of CDH1 expression libraries (Fig S4) were sequenced SE100 by the University of Wisconsin Biotechnology Center on a HiSeq 2500. Sequenced reads were processed using Trimmomatic with the following parameters: LEADING:3 TRAILING:3 CROP:100 HEADCROP:10 SLIDINGWINDOW:4:15 MINLEN:36. Reads were aligned to the mm9 and hg18 genome using RSEM-1.2.4 with a mismatch per seed set to 2 and seed length set to 28 (-bowtie-m 200 --bowtie-n 2 --forward-prob 0.5 --seed-length 28 --paired-end). A matrix of unnormalized reads mapping to the mouse or human genome was generated with R. DE genes were called with EBSeq (Leng *et al*, 2013) using the human matrix for normalization (MedianNorm(human_matrix)). Differentially expressed genes were filtered to have a posterior probability DE greater than 0.95 and a 2 or greater posterior fold change (PostFC). Expression changes reported are PostFC values determined by EBSeq unless otherwise noted to be TPM. Cluster3.0 was used to perform k-means clustering of DE genes (de Hoon *et al*, 2004). Clusters were visualized as Log2FC relative to MEF TPM with Java TreeView (Saldanha, 2004). Motif discovery was performed using HOMER Motif Analysis (Heinz *et al*, 2010) with the following parameters: findMotifs.pl -start −1000 -end 100 -len 6,10. Gene Ontology was performed using HOMER and DAVID (Huang *et al*, 2009). DAVID parameters included: GOTERM_BP_4 and GOTERM_MF_4, viewed by functional annotation clustering. GO Terms within functional annotation clusters with an enrichment score more than 2 were considered significant. DE genes were compared to a published shRNA iterative screen dataset (Borkent *et al*, 2016).

### Dot1L target gene scRNA-seq analysis

Expression of potential Dot1L targets was assessed with single cell RNA-seq data (GEO: GSE108222). Briefly, cells expressing the gene of interest were displayed on a t-distributed Stochastic Neighbor Embedding (t-SNE cluster) plot of MEFs, ESCs, and reprogramming cells in FBS constructed using Monocle2 v2.6.3 on R version 3.4.3 with cells that passed quality control as previously described (Tran *et al*, 2019).

### ChIP-Seq analysis

Data with the accession number GSE90895 (Chronis *et al*, 2017) was downloaded from Gene Expression Omnibus and aligned to mm9 using bowtie2 (Langmead & Salzberg, 2012) with the default parameters. Sam files were converted into Bam files and sorted with samtools-1.2 (Li *et al*, 2009) with the default parameters. Peaks were called relative to the input with MACS2 (Zhang *et al*, 2008) using the following parameters: --broad -p 0.0001. Peak files were annotated using EASeq (Lerdrup *et al*, 2016) from the center of the peak to the nearest gene center. Genes with peaks were visually mapped back on to the Log10 TPM versus Log2 PostFC using R ggplot2 (geom_point graph).

### Constructs

*Brn2* (Addgene 27151), *Ascl1* (Addgene 27150), *Myt1l* (Addgene 27152) were a kind gift from Dr. Marius Wernig. Nanog (Addgene 16578), *Cdh1* (Addgene 71366), and *Myod* (Addgene 8398) were acquired from Addgene. The CDS of *Cdh1* was amplified from Addgene 71366 and moved into the BstBI and NotI sites of pCDH-CMV-MCS-EF1α-Neo. The *Myod* CDS, cut with EcoRI, was moved into the pCDH-CMV-MCS-EF1α EcoRI site.

### Immunofluorescence

Coverslips were fixed in 4% paraformaldehyde-PBS, permeabilized in 0.5% Trition-X-PBS, and washed in 0.2% Tween-20-PBS. Coverslips were blocked for 30 min in blocking buffer (1x PBS, 5% donkey serum, 0.2% Tween-20, and 0.2% fish skin gelatin). Cells were stained for 1 hr with primary antibody in blocking buffer, rinsed 2x in wash buffer, and stained for 1 hr with secondary (1:1000) in blocking buffer. Coverslips were rinsed with wash buffer, stained with DAPI (0.1 μg/ml) in wash buffer, and rinsed with wash buffer. The following antibodies were used for immunofluorescence: anti-murine NANOG (Cosmo Bio RCAB0002P-F, 1:100), anti-human NANOG (R&D Systems AF1997, 1:100), anti-TUJ1 (Novus Biologicals MAB1195, 1:500), anti-CDH1 (eBiosciences 14-3249-80, 1:100) anti-DPPA4 (Thermofisher PA5-47530, 1:100), and anti-DESMIN (BD Biosciences 550626, 1:100). Colony counts and imaging were performed on Nikon Eclipse Ti using NIS Elements software.

### Immunoblot

Whole cells were lysed in RIPA buffer (150 mM NaCl, 1% NP-40, 0.5% Na deoxycholate, 0.1% SDS, 25 mM Tris pH 7.4) with 1x protease inhibitor (Roche 04693116001), sonicated for 5 secs at 20% amplitude, and quantified by cell count or with the DC Protein Assay (BioRad 5000112) according to the manufacturer’s instructions. Equal amounts of protein were loaded onto an SDS-Page gel and transferred to a nitrocellulose membrane. Membranes were blocked in blocking buffer (5% milk, 0.1% Tween-20, 1x PBS) followed by incubation with primary antibody in blocking buffer. Membranes were washed 0.1% Tween-20-PBS and incubated with secondary antibody in blocking buffer. Membranes were washed and visualized with ECL reagent. Images were quantified using Image Studio Lite software. Primary antibodies included: anti-H3K79me2 (Abcam ab3594, 1:1000), anti-Histone 3 (Abcam ab1791, 1:3000), and anti-α-TUBULIN (Cell Signaling 3873, 1:5000).

### Flow cytometry and sorting

For cell cycle determination, cells were fixed in 1 volume of PBS with 9 volume of cold 70% ethanol and permeabilized by freezing at −20 for at least 2 hours. Cells were washed with FACS buffer (1xPBS, 2% FBS, 1mM EDTA), and stained for 30 min in 100 μL FACS buffer containing 1 μL α-KI-67 per million cells, as previously described (Kim & Sederstrom, 2015). Cells were washed with FACS buffer and DNA was stained for 20 min with 50 μg/ml of propidium iodide (in 1xPBS and 2 mM MgCl_2_) supplemented with 100 μg/ml RNaseA. Live cells were used for surface marker staining. Cells were washed in 1x PBS and incubated with antibodies in 1xPBS with 1% FBS for 1 hour. Sorting was performed on a BD FACS AriaII (UW Carbone Cancer Center, Grant #: 1S10RR025483-01) with appropriate controls for gating. FACS quantitation was performed on a BD Accuri C6 Flow Cytometer. Primary antibodies used were anti-THY1-PE (eBioscience 12-0903-81, 1:100), anti-CDH1-eFluor660 (eBioscience 50-3249-80, 1:100), and anti-KI-67-Alexa Fluor 488 (BioLegend 151204).

### siRNA transfection

MEFs were plated at a confluency of 20,000 cells per 24 well on coverslips. Cells were transfected 24 hours after plating with 20 nM siRNA using DharmaFECT1 (Dharmacon T200104) according to the manufacturer’s instructions. Reprogramming was initiated immediately using 2 μg/ml of doxycycline with either vehicle (DMSO) control, 5 μM SGC0946 (ApexBio A4167). Cells were transfected every two days during reprogramming, and siRNA was increased to 40 nM at day 4 to account for increased cell number. Knockdown efficiencies were determined 48h after the second transfection. siHoxd12 (M-046274-01), siNfix (MQ-045912-01), siFosl1(MQ-040704-01) were purchased from Dharmacon. siHic1 (mm.Ri.Hic1.13.1), siMeox2 (mm.Ri.Meox2.13.1), siTwist2 (mm.Ri.Twist2.13.1,) were purchased from IDT.

### RT-qPCR

RNA was isolated using the Isolate II RNA Mini Kit (Bioline BIO-52072), and 1 μg was converted to cDNA using the qScript cDNA Synthesis Kit (Quantabio). Technical replicates of 20 ng were used to measure Ct on a BioRad CFX96 thermocycler with iTaq UniverSYBR Green SMX (BioRad 1725125). Relative expression was calculated against the mean of two housekeeping genes. Primers used in this study:

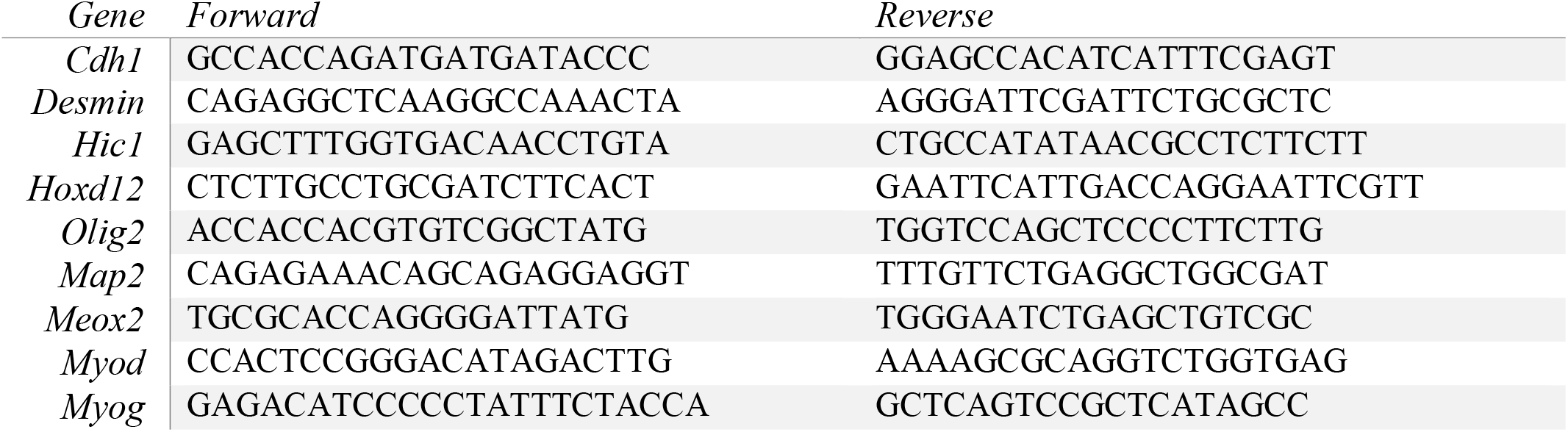
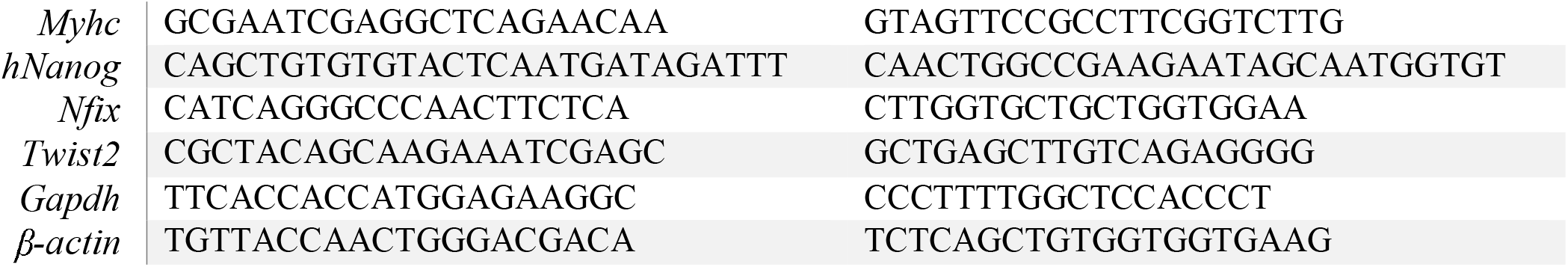

### Statistical analysis

Significance was calculated using the two-tailed t-test function in Graphpad Prism. All experimental replicate information is listed in each figure legend.

## ACKNOWLEDGEMENTS

This work was supported by a UW-Madison Stem Cell and Regenerative Medicine Center postdoctoral award to C.K.W., and Shaw Scientist award to the R.S. lab. We thank Dr. Marius Wernig for the transdifferentiation vectors, Dr. Jayshree Samanta for the TUJ1 antibody, Dr. Roice Wille for R script design, Stefan Pietrzak and Brenton Halvorson for scRNA-Seq analysis, and the R.S. lab members for critical reading of the manuscript.

