## Supplemental figures for "Dot1L methyltransferase activity is a barrier to acquisition of pluripotency but not transdifferentiation"

### Figure S1

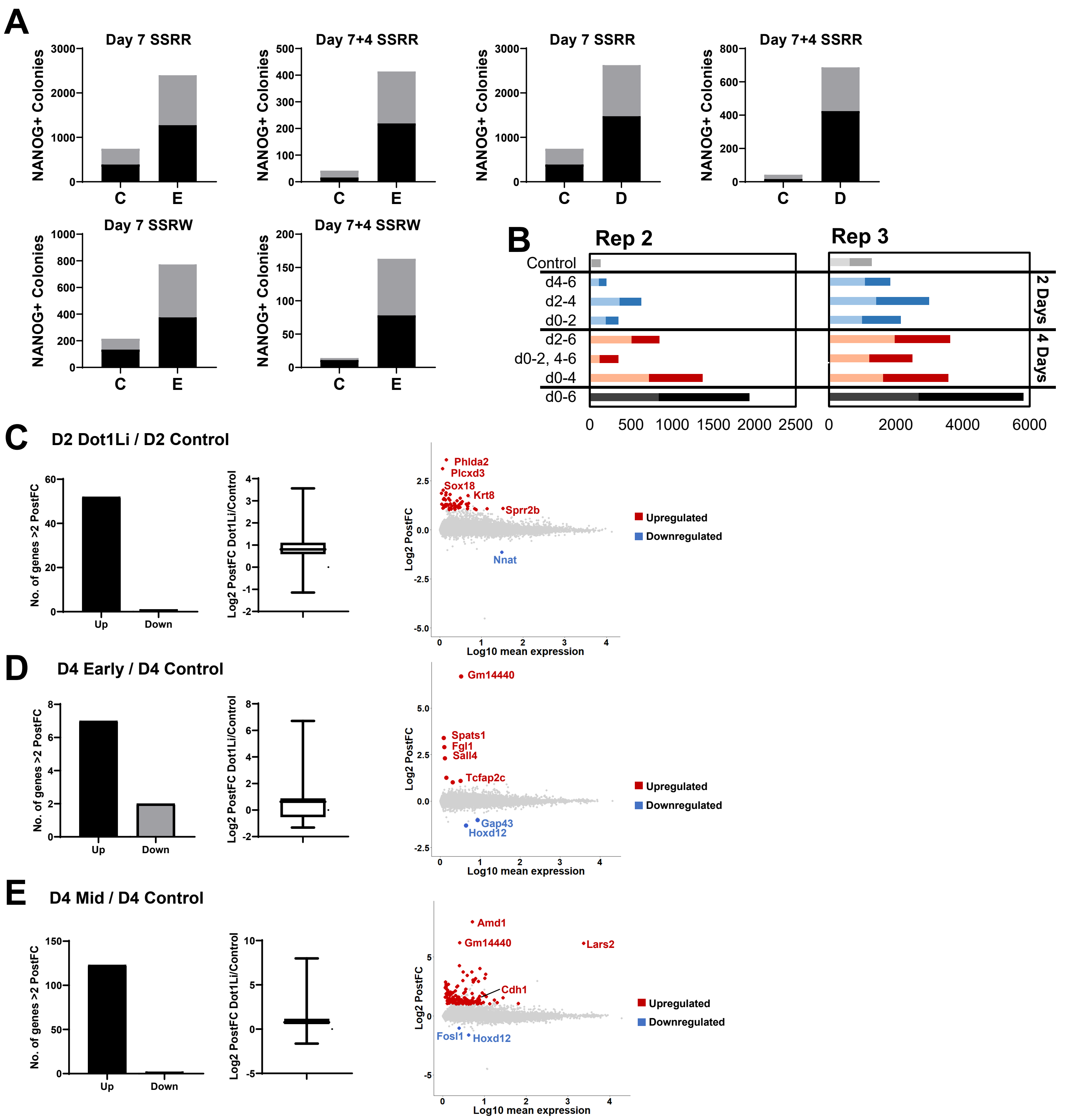


**Figure S1. H3K79me is a barrier for MEF reprogramming with few transcriptional effects.**

A. Number of NANOG+ colonies on day 7 of reprograming and stable NANOG+ colonies 4 days post dox withdrawal of MEFs treated with control (C), Dot1Li (D) or EPZ5676 (E). SSRW and SSRR indicates genotype of MEFs. S = Stemcca cassette of reprogramming factors; R = reverse tetracycline transactivator.

B. Two independent biological replicates of Fig 1C, each comprised of two technical replicates (stacked) of NANOG+ colonies on day 6 of reprogramming.

C-E. Left: Number of genes upregulated or downregulated more than 2-fold (PostFC) with a posterior probability of differential expression greater than 0.95 determined by EBSeq. Middle: Box plot of Log2 PostFC of all genes with a posterior probability of differential expression greater than 0.95. Right: Log10 average TPM of the two samples versus Log2 PostFC of all genes. More than 2-fold upregulated indicated in red and downregulated indicated in blue. C. Day 2 Dot1Li versus day 2 Control, D. Day 4 Early (days 0-2) Dot1Li versus day 4 Control, E. Day 4 Mid (days 2-4) Dot1Li versus day 4 Control

#### Figure S2

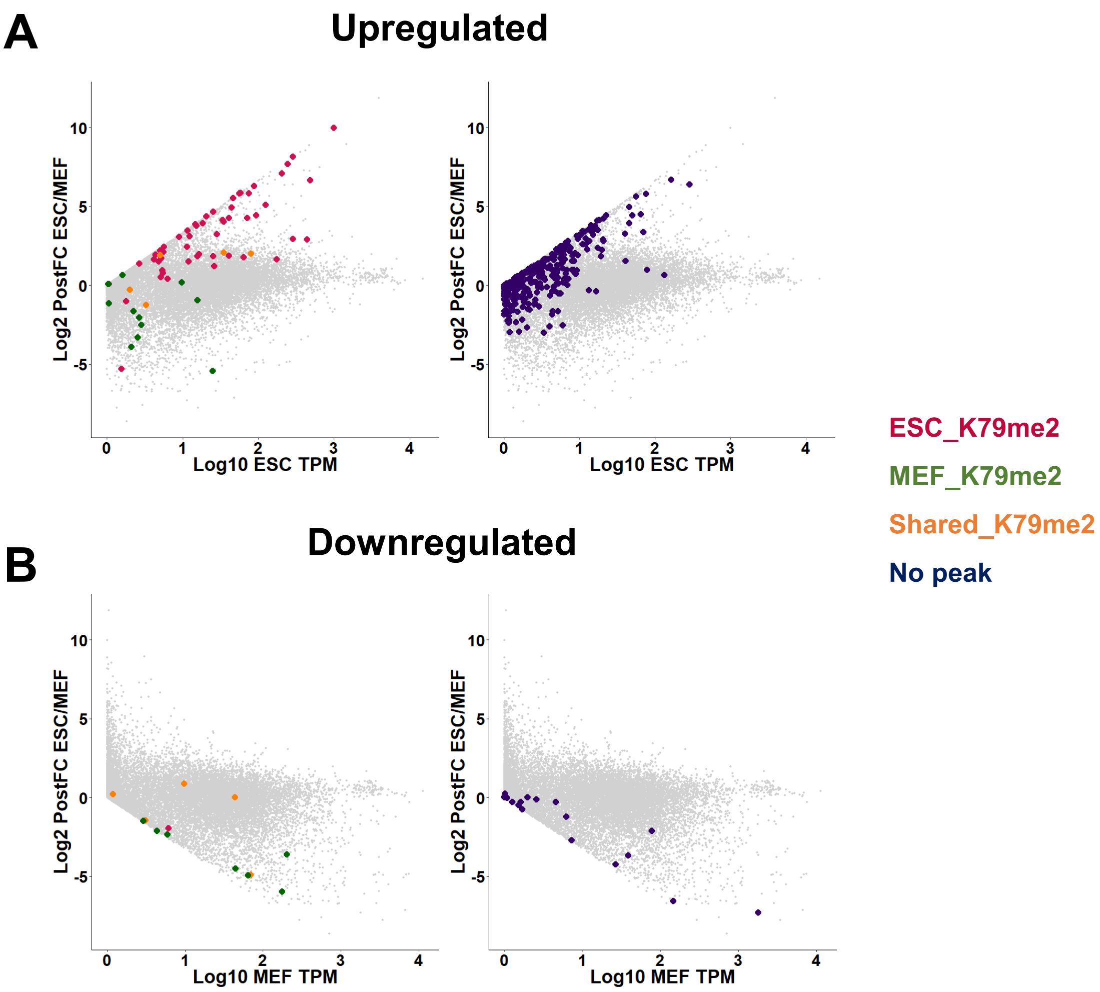


**Figure S2. H3K79me2 is enriched on numerous genes yet few change transcriptionally.**

A.H3K79me2 peak status of Dot1Li upregulated genes designated by color, plotted on Log10 ESC TPM versus Log2 PostFC in ESCs relative to MEFs of all genes.

B. H3K79me2 peak status of Dot1Li downgregulated genes designated by color, plotted on Log10 MEF TPM versus Log2 PostFC in ESCs relative to MEFs of all genes.

### Figure S3

## A

| Motif | Cluster | p-value | Percent of targets | Function |
|---|---|---|---|---|
| **E-Box (SNAI2) (ZEB1)** | II<br>III | 1E-02<br>1E-10 | 50.0%<br>39.8% | Repressor |
| **TCF7** | IV | 1E-7 | 16.7% | Activator |
| **HOXD12** | V | 1E-05 | 2.1% | Activator |
| **HIC1** | II/III<br>V | 1E-05<br>1E-02 | 4.0%<br>77.8% | Repressor |

## B

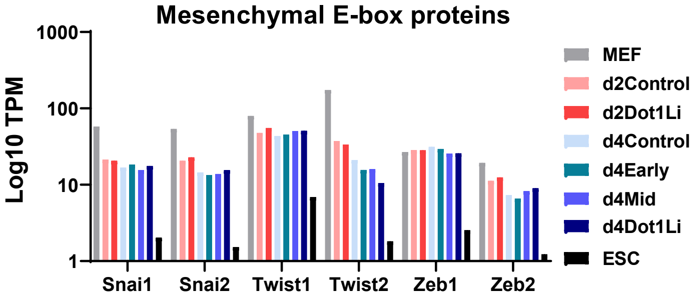


## C

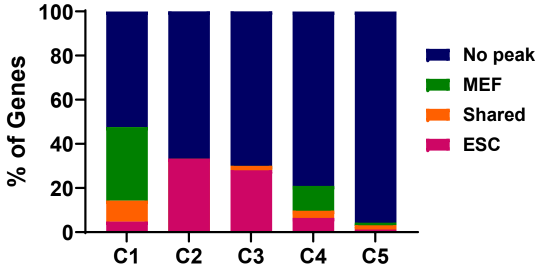


**Figure S3. Dot1L inhibition promotes aberrant transcriptional changes.**

A. Table of motifs in each cluster identified with HOMER. The p-value, percent of targets within the cluster, and function of the binding protein for motifs bound by Dot1Li-DE genes are displayed.

B. Log10 TPM bargraph of mesenchymal E-box binding proteins.

C. Bargraph of the percentage of genes with an H3K79me2 called peak in each cluster.

### Figure S4

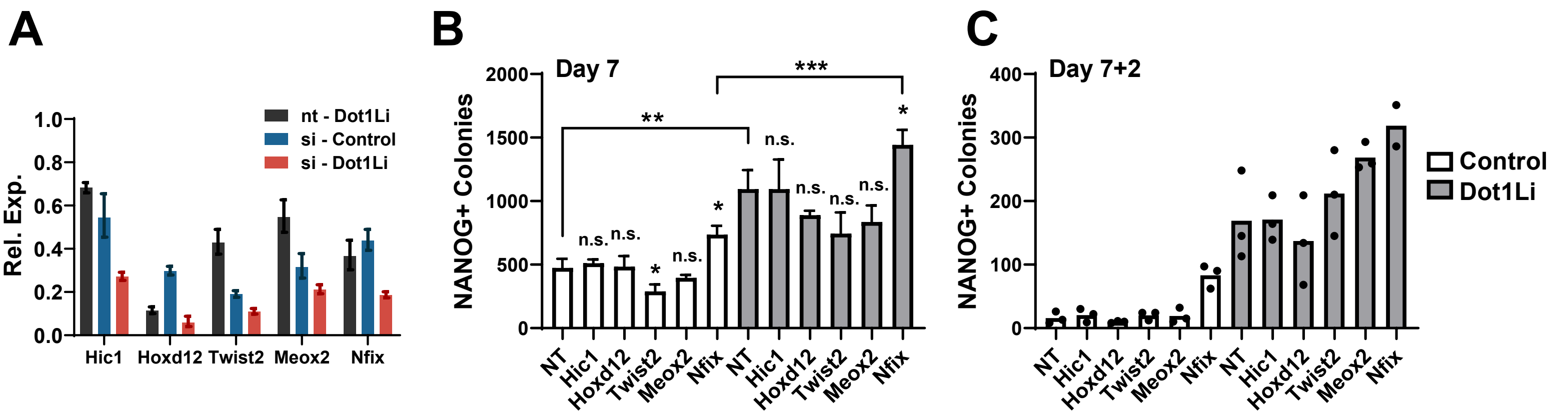


**Figure S4. Inhibition of Dot1L enhances reprogramming beyond modulation of single genes.**

A. Relative expression of si-depleted genes on day 4 of reprogramming. Cells transduced with non-targeting (NT) siRNA treated with control set to 1.

B. An independent biological replicate of Fig 4E, consisting of 3 technical replicates. ***P<0.001, **P<0.01, *P<0.05, and not significant (n.s.) P>0.05 by unpaired t-test.

C. An independent biological replicate of Fig 4F, consisting of 2 or 3 technical replicates (shown as black dots).

### Figure S5

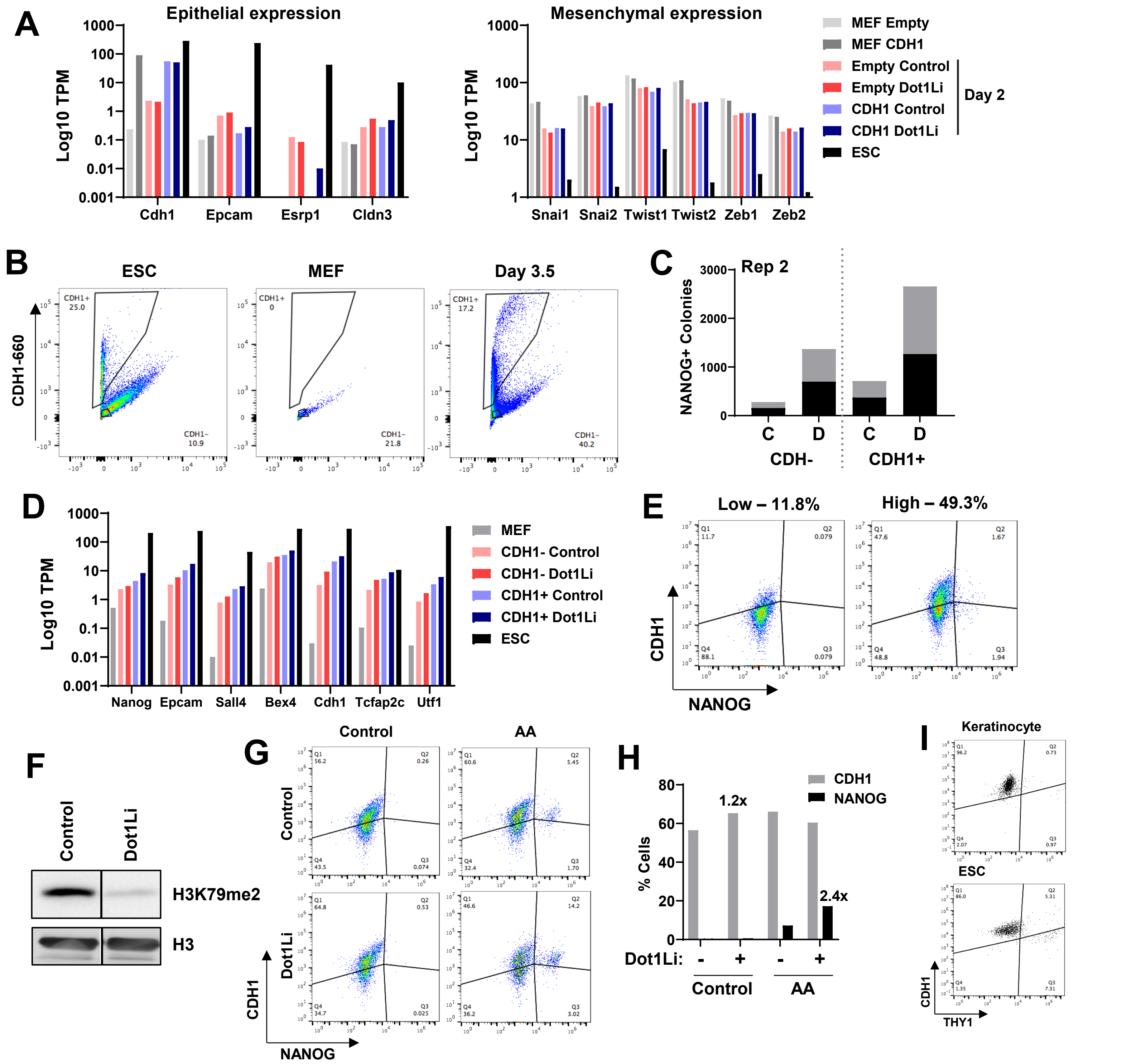


**Figure S5. Inhibition of Dot1L enhances reprogramming of epithelial cells.**

A. Log10 TPM bargraph of MEFs and day 2 reprogramming cells transduced with empty vector control or *Cdh1*, treated with and without Dot1Li.

B. Flow cytometry sorting of CDH1 positive and negative cells for Fig 5C with ESC and MEF controls. Gates indicate collected cells.

C. Biological replicate of Fig 5C comprised of two technical replicates (stacked) of NANOG+ colonies on day 7.5 of reprogramming.

D. Log10 TPM bargraph of representative upregulated genes in CDH1- and CDH1+ that overlapped with genes upregulated in ESCs relative to MEFs (Fig 5E).

E. CDH1 surface expression measured by flow cytometry in two pre-iPSC lines, “low” and “high” CDH1.

F. H3K79me2 and histone 3 (H3) loading control immunoblot on day 4 of pre-iPSC reprogramming with Dot1Li or control.

G. CDH1 and NANOG expression measured by flow cytometry on day 10 of “high CDH1” pre-iPSCs reprogramming with ascorbic acid (AA).

H. Quantification of CDH1 (gray bars) and NANOG (black bars) measured by flow cytometry (Fig S5G).

I. Flow cytometry analysis of THY1 and CDH1 on ESCs and keratinocytes.

### Figure S6

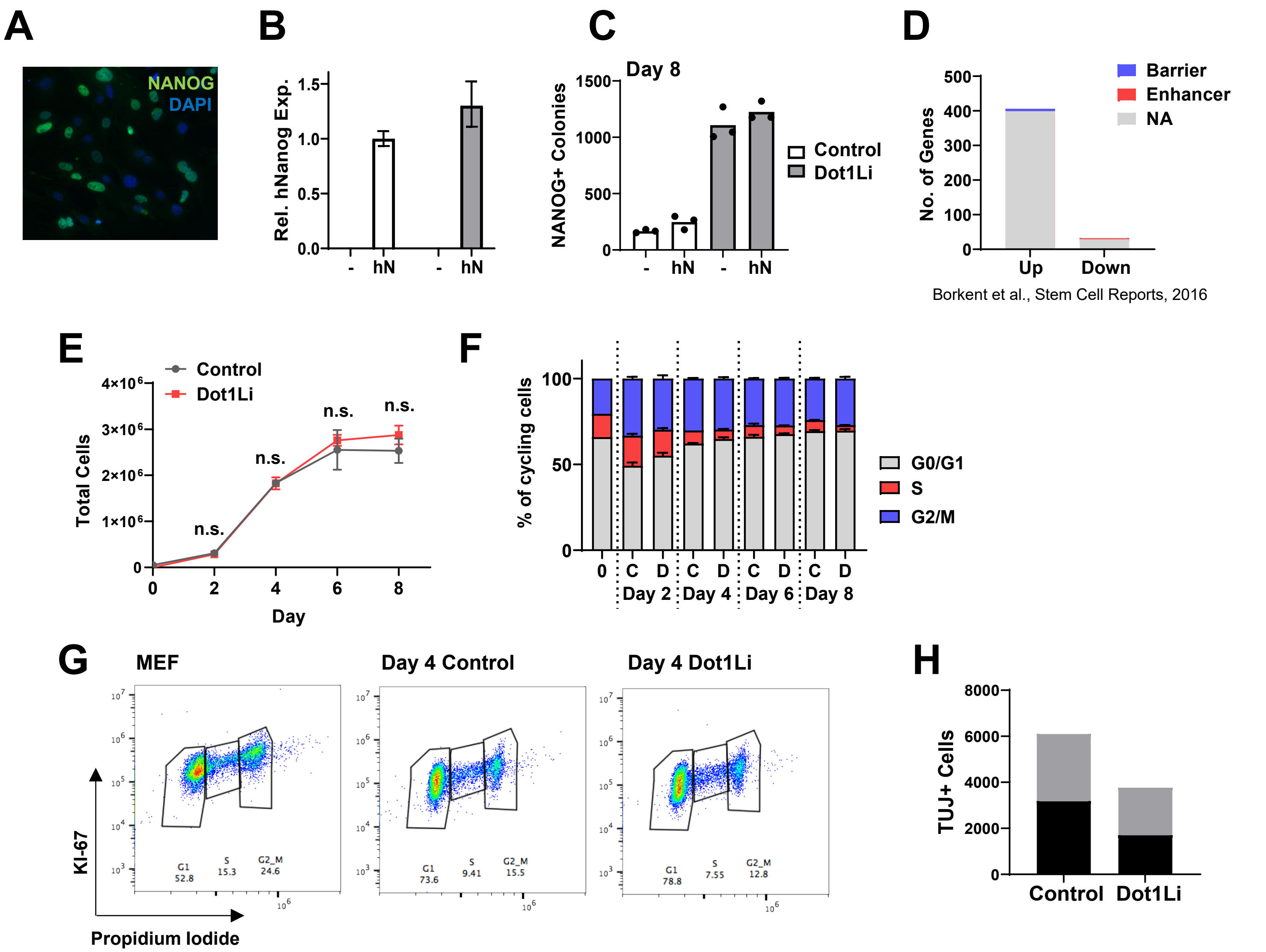


**Figure S6. Dot1L does not maintain cellular identity by regulation of reprogramming factors or cell cycle.**

A. Immunofluorescence of lentivirally transduced human NANOG in MEFs.

B. Relative human *Nanog* expression measured on day 4 of reprogramming in cells treated with control (white bars) or Dot1Li (gray bars). Control treated cells transduced with *hNanog* set to one.

C. An independent biological replicate of Fig 5I consisting of 3 technical replicates (shown as black dots).

D. Overlap of Dot1Li-DE genes and Borket *et al,* 2016 screen. Genes chosen for overlap affected reprogramming more than 2-fold. “Barriers” indicate genes targeted by shRNAs enriched in reprogrammed cells and “enhancers” are genes targeted by shRNAs depleted in reprogrammed cells.

E. Cells treated with control (gray) or Dot1Li (red) were counted every two days during reprogramming. Error bars represent the standard deviation of three technical replicates. Not significant (n.s.) P>0.05 by unpaired t test.

F. Quantification of cell cycle analysis. Cells on day 0 (0), and cells treated with Dot1Li (D) or control (C) were assessed every two days during reprogramming.

G. Representative propidium iodide and KI-67 flow cytometry cell cycle analysis (Fig S6F).

H. Independent biological replicate of Fig 6B comprised of two technical replicates (stacked) of TUJ+ neurons on day 7 of transdifferentiation.
